# The structure of a human translation initiation complex reveals two independent roles for the helicase eIF4A

**DOI:** 10.1101/2022.12.07.519490

**Authors:** Jailson Brito Querido, Masaaki Sokabe, Irene Díaz-López, Yuliya Gordiyenko, Christopher S. Fraser, V. Ramakrishnan

**Affiliations:** MRC Laboratory of Molecular Biology, Cambridge CB2 0QH, United Kingdom; Department of Molecular and Cellular Biology, College of Biological Sciences, University of California, Davis, CA 95616, USA

## Abstract

**Summary:** Initiation of mRNA translation is a key regulatory step in gene expression in all eukaryotes. Canonical initiation of translation in eukaryotes involves recruitment of the 43S preinitiation complex to the 5′ end of mRNA by the cap-binding complex eIF4F to form the 48S initiation complex (48S), followed by scanning along the mRNA until the start codon is selected.1–8 We have previously shown that eIF4F binds near the mRNA channel exit site of the 43S, leaving an open question about how mRNA secondary structure is removed as it enters the mRNA binding channel on the other side of the 40S subunit.^4^ Here we describe a human 48S positioned at the start codon that shows that in addition to the eIF4A that is part of eIF4F, there is a second eIF4A helicase bound to the mRNA entry site. The entry channel bound eIF4A is positioned through interactions with eIF3 and the 40S subunit to enable its ATP-dependent helicase activity to directly unwind secondary structure located downstream of the scanning 48S complex. The structure also reveals universally conserved interactions between eIF4F and the 48S, likely explaining how this complex can promote mRNA recruitment in all eukaryotes. mRNA translation has emerged as an important tool for developing innovative therapies, yet several fundamental aspects of its regulation remain unknown. This work sheds light on the critical regulatory roles of eIF4A and eIF4F during the recruitment and scanning of the 5′ UTR of mRNA.

## Introduction

Initiation of translation in eukaryotes involves over 20 different initiation factors (eIFs). The process starts with the assembly of the 43S preinitiation complex (43S) consisting of the 40S ribosomal subunit bound to initiation factors eIF1, eIF1A, eIF3 and eIF5 and a ternary complex (TC) of eIF2, guanosine 5′-triphosphate (GTP), and methionyl initiator transfer RNA (tRNAiMet). In parallel, the cap-binding complex eIF4F bound to the 5′ end of mRNA recruits the 43S to form the 48S translation initiation complex, which then scans along the mRNA until it encounters a start codon.1–8 The cap-binding complex eIF4F consists of a scaffold protein eIF4G, an m^7^G cap-binding protein eIF4E, and a DEAD-box helicase eIF4A.^9^ Additionally, eIF4G interacts with poly(A)-binding protein PABP located at the 3′ untranslated region (UTR) of mRNA, which further brings the 3′ and 5′ ends close together. In metazoans, the attachment of the 43S to the mRNA-eIF4F-PABP complex is mediated by a direct interaction between eIF3 and eIF4G.3–5,8,10,11 Recently, we determined the structure of a human 48S complex during scanning, which revealed the binding site of eIF4F.^4^ While the resolution was modest, the structure showed that eIF4F interacts with eIF3e and eIF3k/l located upstream of the 43S, near the mRNA channel exit site. This finding was consistent with a slotting mechanism of mRNA recruitment and explained the presence of a blind spot between the m^7^G cap structure and the recognition of an initiation codon. Nevertheless, the location of eIF4F in the 48S left an unanswered question about how eIF4A that is part of eIF4F behind the scanning 40S subunit could act as a helicase during scanning and actively unwind secondary structure before it enters the mRNA binding channel.

The helicase activity of eIF4A is stimulated by eIF4G and eIF4B.12–14 In addition, S. cerevisiae eIF4B likely induces conformational changes in the 40S subunit to facilitate the attachment of the 43S to mRNA. Yet, it is unclear how eIF4B interacts with eIF4A and/or eIF4F on the surface of the 40S subunit. Interestingly, the ATPase activity of yeast eIF4A does not appear to require eIF4G and eIF4E15, whereas in mammals, recruitment and scanning requires the cap-binding complex eIF4F.^13^ Thus, it remained uncertain how eIF4A, eIF4B, and the eIF4F complex could work together to unwind secondary structure during mRNA recruitment and scanning.

In our prior structure, we used an mRNA that lacked an AUG codon and was shorter than the footprint of the 48S predicted by the structure^4^. Here we capture a later stage initiation complex of the 48S complex positioned over a start codon, using a longer more physiological mRNA with a long and structured 5′ UTR, AUG codon, and a 3′ UTR ending in a poly(A) tail. Unexpectedly, a second eIF4A helicase is bound at the mRNA entry channel of the 48S complex. This eIF4A helicase interacts with universally conserved eIF3 subunits and is entirely separate from the eIF4F complex positioned at the mRNA exit site on the other side of the 40S subunit. The structure identifies the interaction of eIF4B with the entry site bound eIF4A, providing insight into its role in regulating this second eIF4A rather than eIF4F during scanning. The discovery of a second eIF4A molecule positioned at the point of entry of mRNA into the mRNA binding channel of the 40S subunit resolves many seemingly contradictory data, including the unanswered question of how secondary structure is unwound by the helicase activity of eIF4A during scanning.

### Structure a human 48S translation initiation complex

To reconstitute as complete and physiological a 48S complex as possible for cryo-EM analysis, we use a capped mRNA containing a long 5′ UTR (105 nucleotides), an AUG codon, followed by a short CDS (9 nucleotides) and a poly(A) tail (∼ 90 nucleotides) (Fig. S1). To promote the proposed interaction of the poly(A) tail with the 5′ end of mRNA during complex formation, we included PABP in addition to a full complement of initiation factors, including full-length eIF4G1. Inclusion of a start codon enables us to generate a late-stage initiation intermediate representing the 48S with the start codon in the P site. To stabilize the complex further, we used Rocaglamide A (RocA), which clamps eIF4A onto polypurine sequences16–19, while also including three polypurine motifs (GA)_6_ in the 5′ UTR of the mRNA.

Single-particle reconstruction using cryo-EM reveals density corresponding to the cap-binding complex eIF4F in the same location as seen in our previous structure (Fig.1, Fig. S2).^4^ Masked classification on the mRNA channel entry site followed by additional masked classification on the region we previously saw for eIF4F yielded a cryo-EM map at 3.1 Å resolution (Fig. S3A). Further masked classification on TC yielded a map with a reduced overall resolution of 3.7 Å (Fig. S3B) but improved the density for TC.

**Figure 1:**
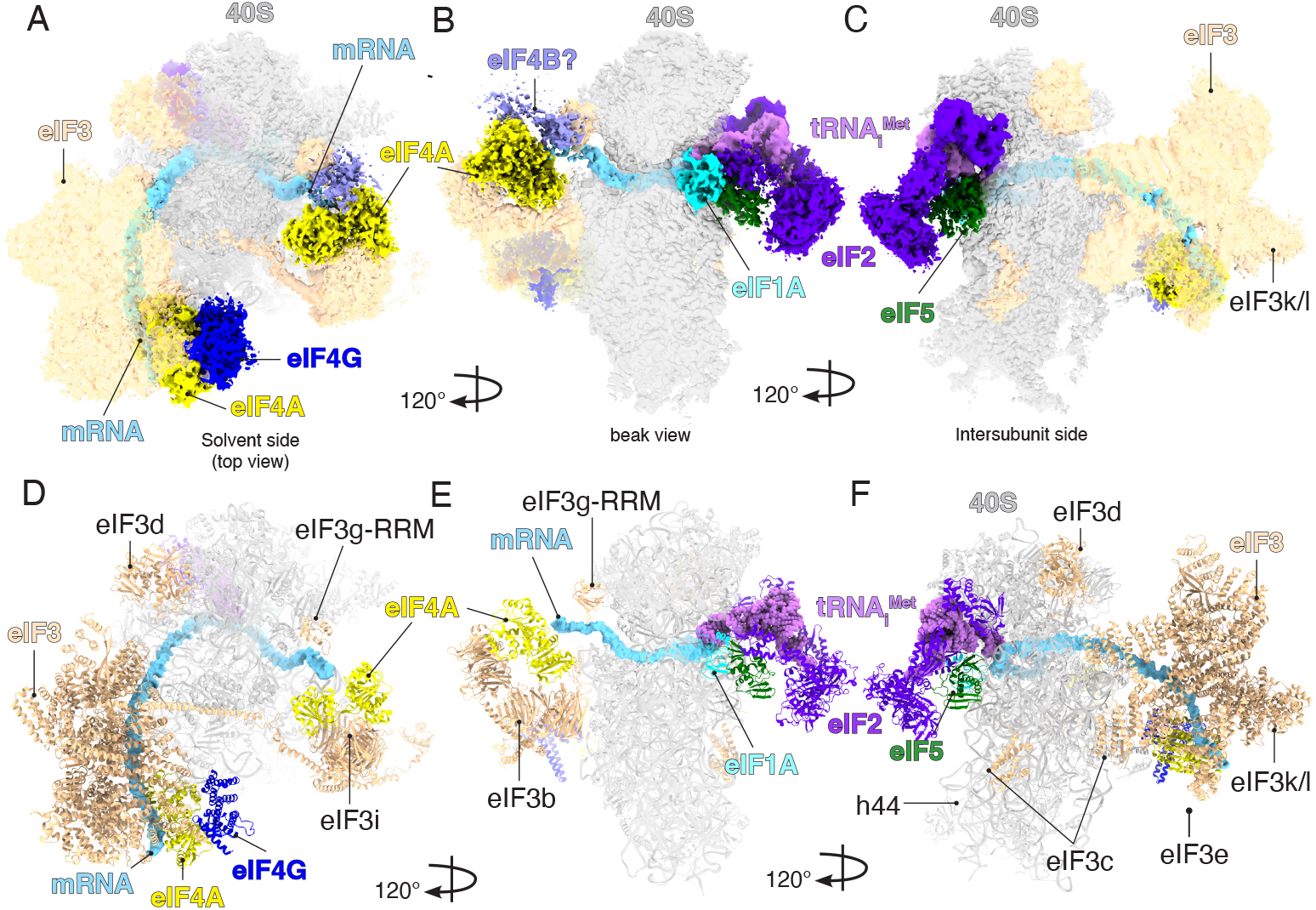
Cryo-EM 3D reconstruction and the structure of human 48S complex. Overview of the cryoEM density map (A to C) and molecular model of human 48S (D-F) shown in different orientations. The map contains densities for 40S small ribosomal subunit, eIF3, eIF1A, eIF5-NTD, eIF2, tRNA_i_^Met^, mRNA, eIF4F (eIF4G and eIF4A), and the second molecule of eIF4A bound to eIF3 at the mRNA channel entry site. The map also contains a possible density for eIF4B in contact with the second molecule of eIF4A at the entry site.

We observe additional density extending out from the eIF3bgi module at the mRNA channel entry site (Fig.1). This density was not present in any previous structures of 48S complexes. The local resolution and detailed shape indicate that this additional density is due to a second eIF4A helicase bound to mRNA, in addition to the eIF4A that is part of eIF4F on the other side of the 40S subunit (Fig. 1). To test whether the presence of this second, entry-site eIF4A is a result of its being trapped by RocA, we determined the structure of a complex without RocA and found that both eIF4F and the second eIF4A are still present (Fig. S4). To assemble this complex, we used a capped mRNA with the 5′ UTR of b-globin (55 nucleotides) followed by a CDS (124 nucleotides) containing a GC-rich region located 27 nucleotides downstream of the AUG. This GC-rich region forms a downstream loop (DLP) in the mRNA and has been used to trap eIF4A in the 48S.^20^

The local resolution of 5.9 Å for the cap-binding complex eIF4F (Fig. S4C and SD) is greatly improved compared to our previous structure^4^, and allows us to see direct interactions not only to additional subunits of eIF3 but unexpectedly to the 40S small ribosomal subunit. The improved density shows secondary structure elements and enables us to accurately model the middle domain of eIF4G (HEAT-1 domain) and eIF4A (Fig. 1). Although the mRNA has an m^7^G cap and a poly(A) tail, we did not see any additional density that could be assigned to eIF4E or PABP.

The overall conformation of the structure is a post-scanning intermediate — the initiator tRNA is inserted completely into the P site (a Pin state) where it bases pairs with the start codon, and the 40S subunit mRNA binding channel is in the closed conformation. Additionally, the overall resolution allowed us to identify density corresponding to the 40S, eIF1A, eIF2a, eIF2b, eIF2g, tRNAiMet, the octameric structural core of eIF3 as well as its peripheral subunits (b, d, g, i) (Fig. 1, Table S1).

### A second eIF4A helicase in the 48S

Compared to our previous 48S structure, we observed additional density at the entry site of the mRNA channel (Fig. 2A-B). Much of this density can be accounted for by eIF4A, which after rigid-body fitting of the known crystal structure of human eIF4A17 shows close agreement with the density, especially the eIF4A-CTD, for which almost the entire secondary structure was resolved (Fig. 2B). eIF4A binds to a pocket formed by eIF3bgi, eIF3a-CTD and rRNA h16 (Fig. 2C-D). This position of eIF4A agrees well with previous biochemical, cross-link mass spectrometry, and proximity labelling data.4,15,21 Moreover, recent singlemolecule data revealed FRET between eIF4A and ribosomal uS1922; the latter is located within FRET distance of this second eIF4A in our structure.

**Figure 2:**
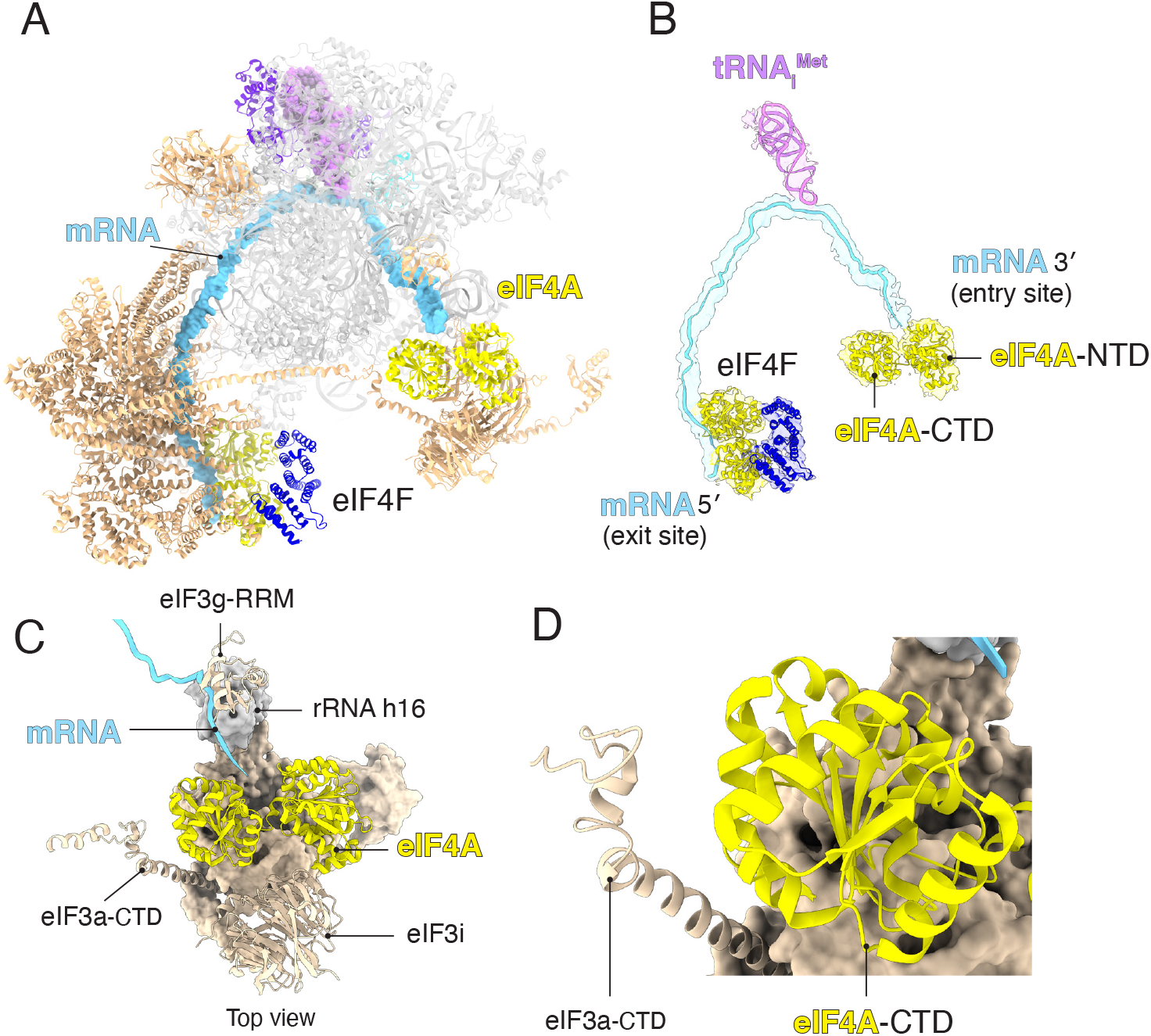
Structure of human 48S reveals a second molecule of eIF4A bound at the mRNA channel entry site. (A-B) Structure of human 48S revealing the path of the mRNA from eIF4F towards the second molecule of eIF4A at the entry site. (B) Atomic model of eIF4A, eIF4F, mRNA, and tRNA_i_^Met^ fitted into the cryo-EM map. (C) Interactions of eIF4A at the entry site. (D) The interface between eIF4A-CTD and eIF3a-CTD.

The two RecA domains of the entry site eIF4A adopt an open conformation that differs from that described for the yeast eIF4A-eIF4G complex^23^ (Fig. S5). Instead, the conformation of this entry site eIF4A resembles that of the crystallographic structure of the eIF4A-Pdcd4 (Programmed cell death 4) complex^24^ (Fig. S5). This conformation allows the NTD and CTD to contact the mRNA while the nucleotide-binding domain adopts a more open conformation. The structure shows that eIF4A could remain bound to mRNA as it cycles through the open and closed conformations during scanning, which may help explain the factor-dependent processivity of human eIF4A described previously.^14^

Unlike previous 48S structures in which mRNA outside the 40S subunit was not seen, there is clear and continuous density for the mRNA which allowed us to trace its path for the first time from the cap-binding complex eIF4F upstream of the 43S, through the 40S subunit mRNA channel all the way to the second molecule of eIF4A downstream of the 43S (40 nt upstream and 20 nt downstream of the AUG; Fig. 2A-B). This allows us to clearly see the nature of the interactions of mRNA with the two eIF4A molecules on opposite sides of the 40S subunit.

It has been shown that yeast 43S stimulates the ATPase activity of eIF4A.^15^ Interestingly, this activity of yeast eIF4A does not require eIF4G or eIF4E. Instead, eIF3, especially eIF3i and eIF3g, are required to stimulate the ATPase activity of eIF4A. Because the cap-binding complex eIF4F binds upstream of the 43S, it was unclear how eIF3 subunits apparently downstream of the 43S could affect the activity of eIF4A. However, it has been shown that the eIF3bgi complex is dynamic, binding to both the mRNA entry channel and the subunit interface.[25]. Thus, it was not clear whether the eIF3bgi domain stimulates the eIF4A helicase directly or indirectly on the 40S subunit surface. This is now solved by revealing the direct interaction between the second molecule of eIF4A and the eIF3bgi module (Fig. 2). Indeed, the structure reveals that eIF3i is the main binding partner of this second eIF4A.

eIF3a binds to the solvent exposed site of the 40S, near the mRNA channel exit site. However, its C-terminal domain extends toward the solvent side of the 40S, where it interacts with eIF3bgi at the mRNA channel entry site.26 Our previous cross-linking mass spectrometry indicated that eIF3a-CTD (Lys632) is in close proximity to eIF4A-CTD (Lys291).^4^ Consistent with this observation, the structure shows a close proximity between eIF3a630-646, a disordered region in the eIF3a-CTD and Lys291 of eIF4A-CTD (Fig. 2C-D). Indeed, the structure also explains prior biochemical data indicating that eIF3a-CTD and eIF3bgi play important role during recruitment and scanning.27.

Mammalian eIF4G has two eIF4A-binding domains, one located in the middle domain (HEAT-1) and a second one located at the C-terminal (HEAT-2).28,29 The eIF4G HEAT-1 is highly conserved amongst eukaryotes and is the core of the eIF4F complex, while the eIF4G HEAT-2, not present in *S. cerevisiae*, is poorly conserved and is not essential.28,29 It has been proposed that HEAT-2 plays a stimulatory role28. To test whether HEAT-2 is required for binding of the second molecule of eIF4A, we analyzed a 48S complex with a truncated eIF4G lacking HEAT-2. The second molecule of eIF4A is still present in this complex (Fig. S6), suggesting that its binding at the mRNA entry site does not require eIF4G, consistent with previous biochemical data showing that *S. cerevisiae* 43S stimulates the ATPase activity of eIF4A in the absence of eIF4G.^15^

A previous biochemical study proposed that eIF4F binds at the mRNA entry site.^8^ However, at the entry site, the domain of eIF4A that interacts with eIF4G in the eIF4F complex is involved in interactions with eIF3i (Fig. 3A-B). The structure suggests that the position of eIF4A at the mRNA channel entry site is incompatible with its interaction with the eIF4G HEAT-1 seen in the cap-binding complex eIF4F (Fig. 3A-B). We therefore propose that this second molecule of eIF4A acts independently of eIF4F.

**Figure 3:**
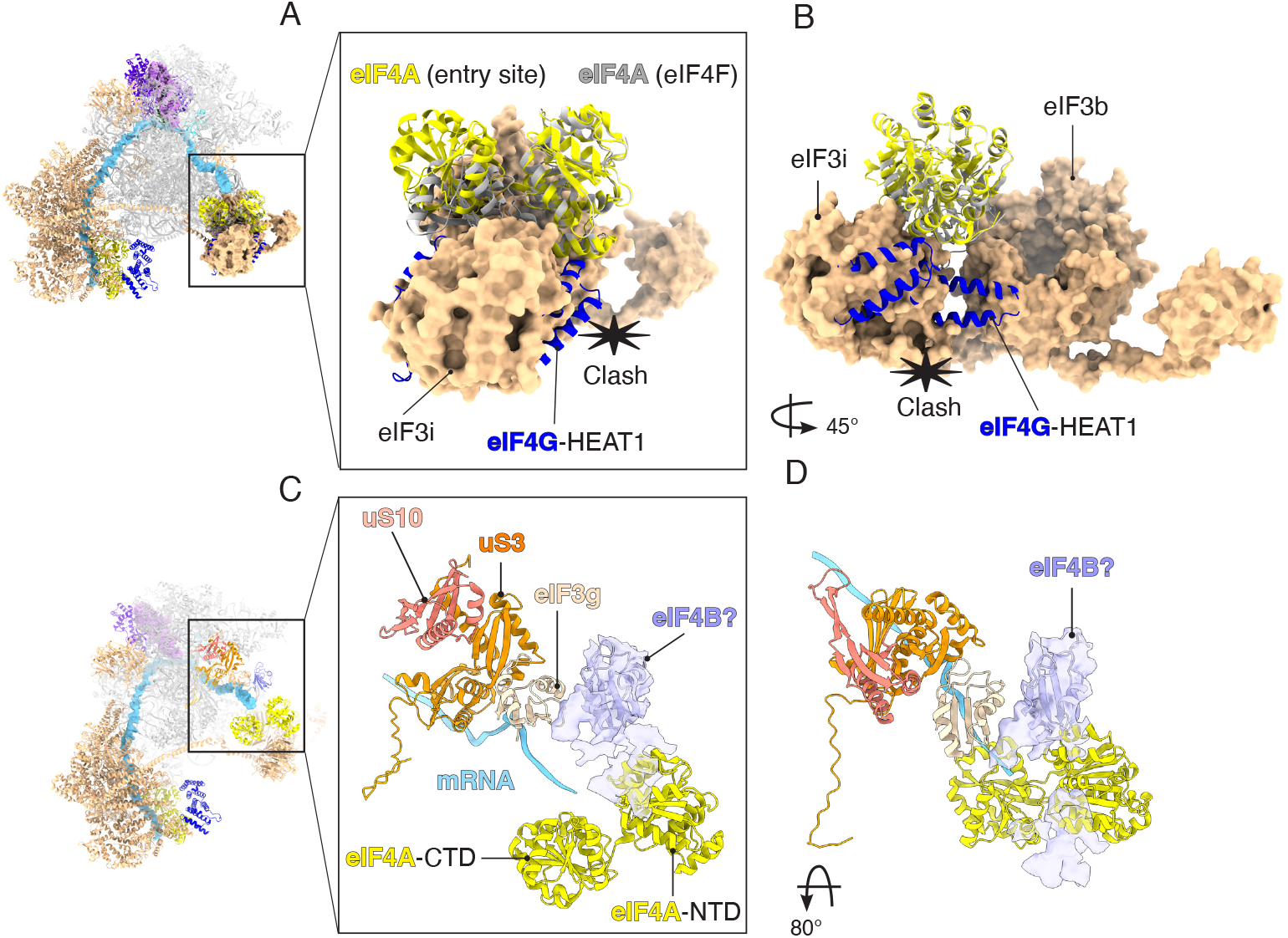
Conformational differences between the two molecules of eIF4A and a possible location of eIF4B in the 48S. (A to B) Superposition of eIF4F with entry site eIF4A to highlight conformational differences. eIF4G-HEAT1 would clash with eIF3i and -b. (C to D) Atomic model of human eIF4B-RRM (AlphaFold prediction) fitted into the unassigned cryo-EM density map at the entry site. The possible location of eIF4B places it in close proximity to eIF4A-NTD, eIF3g-RRM, uS3, and uS10.

### A possible location for eIF4B

By itself, eIF4A has low helicase activity. Even though its activity is modulated by the 43S^15^, the helicase activity of eIF4A in scanning the 5′ UTR of mRNA with stable secondary structure is enhanced by the cofactor eIF4B or eIF4H.12,13The precise binding site of eIF4B on the 48S remained elusive. Our cryo-EM map contains an unassigned density adjacent to the eIF4A-NTD, in close contact with eIF3g-RNA recognition motif (RRM) and ribosomal protein uS10 (Fig 3C-D). Although the resolution of the additional density is low, its size and shape are consistent with the eIF4B-RRM (Fig. 3C-D). Consistent with this interpretation, cryo-EM maps of complexes formed without eIF4B does not contain this additional density (Fig. S7). This location of eIF4B agrees well with previous biochemical data indicating that eIF4B or eIF4H interact with the same region of the eIF4A-NTD.30 Furthermore, previous yeast two-hybrid analysis identifying the ribosomal protein uS10, located in close proximity to the unassigned density, to be the main interaction partner of eIF4B.^12^ This possible location, which suggests a direct interaction between eIF4B and the entry-site eIF4A, agrees well with the role of eIF4B in increasing the directionality of eIF4A translocation.^14^

In addition to its role as a subunit of eIF4F, we propose that a second molecule of eIF4A at the mRNA channel entry site likely promotes mRNA recruitment and scanning. A second eIF4A in the 48S is also consistent with the finding that eIF4A is in molar excess over other components of eIF4F.31,32 Nevertheless, even a second eIF4A in the 48S does not account for the substantial excess eIF4A compared to eIF4F in the cell.31,32 Thus, it possible that eIF4A could have additional functions apart from its role in the 48S complex. Consistent with this idea, the eIF4A-mRNA interaction has a long lifetime^33^, and eIF4A is known to be able to melt RNA secondary structure in the absence of the 43S.31

### eIF4F interacts with eIF3 and ribosomal protein eS7

The recruitment of the 43S to the 5′ UTR of mRNA is a critical step in the translation of eukaryotic mRNAs. Our previous structure of a human 48S complex allowed us to identify the location of eIF4F^4^, but the low local resolution and the adjacent highly flexible region of the 43S meant that we could only infer interactions with non-core subunits of eIF3 (eIF3e, -k, -l). Since these subunits do not exist in *S. cerevisiae*, it was also not clear how eIF4F could interact with the yeast 43S complex. The improved local resolution in this region allows us not only to build eIF4F more accurately but also identify its interactions with the 43S in far greater detail, including additional ones that are likely universally conserved. The density shows almost the entire secondary structure of the middle domain of eIF4G (HEAT-1) and eIF4A (Fig. 4A-C). Rigidbody fitting of a crystal structure of a human eIF4A (PDB:5ZC9)^17^ allowed us to assign its corresponding density (Fig. 4D-F). A model of the middle domain of eIF4G based on an AlphaFold prediction^35^ was used for rigid-body fitting into the density (Fig. 4D-F). The predicted model agreed well with the density, allowing us to locate the domain of eIF4G in the structure.

**Figure 4:**
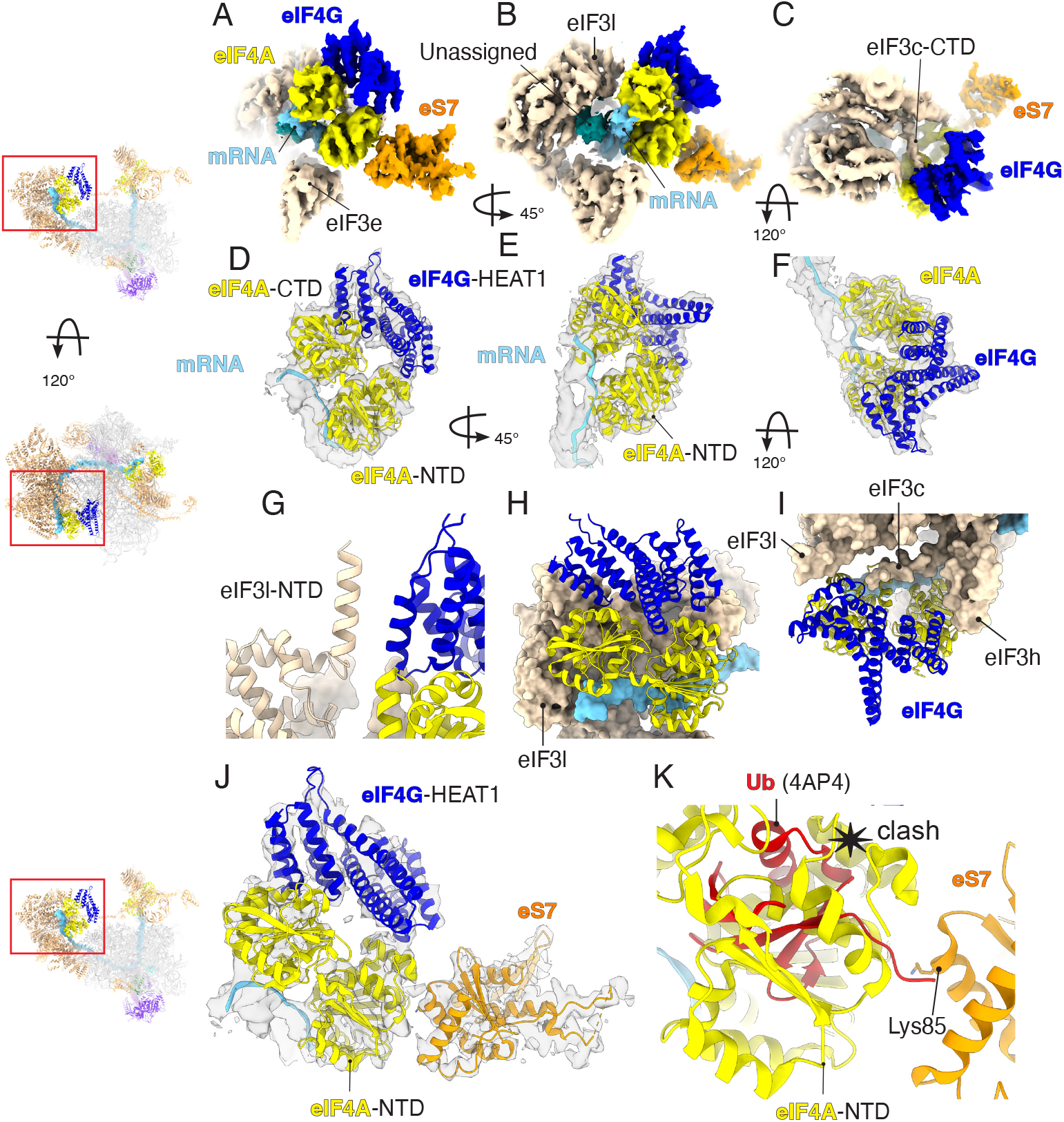
Structure of eIF4F and its interactions with eIF3 and ribosomal protein eS7. (A to C) Cryo-EM map to highlight eIF4F and its interaction network. (D to F) Atomic model of human eIF4A and eIF4G fitted into the cryo-EM map. (G to I) The insets highlight the interaction network of eIF4F with eIF3. (J) Close up to highlight the interaction between eIF4F and ribosomal protein eS7. (K) Superposition of ubiquitin (PDB:4AP4)^34^ with the structure of human 48S to highlight the possible clash between eIF4A and ubiquitin at its predicted binding site (Lys85).

eIF4F is positioned on the 5′ side of mRNA relative to the 40S subunit, near the mRNA channel exit site. For the first time, mRNA interacting with eIF4A is clearly visible (Fig. 4A-C), and eIF4A adopts a structure intermediate between the extended conformation, found in a crystal structure of a yeast eIF4A-eIF4G complex without mRNA in which the two RecA domains are rotated away from each other23, and the closed conformation of an eIF4A-mRNA dimeric complex17. This conformation allows the two RecA domains to interact with the mRNA while maintaining their interactions with eIF4G (Fig. 4A-F and Fig. S8).

In metazoans, eIF3 plays a critical role during the recruitment of 43S to the eIF4F-mRNA.3–5,8,10 The structure reveals an unexpectedly large interaction network between eIF3 and eIF4F (Fig. 4G-I). Previously, because of high flexibility, the structure of eIF3l-NTD remained elusive. The improved local resolution reveals a near-complete structure of eIF3l-NTD and its interaction with eIF4G. Interestingly, the interaction between eIF3l-NTD and eIF4G occurs through a domain of eIF4G located between residues Met775 and Ala787, which was unexpected (Fig. 4G). Additionally, the cryo-EM map reveals an unassigned density connecting eIF4G with eIF3c and -h (Fig. 4). Although the local resolution does not allow us to assign this density unambiguously, it likely belongs to the C-terminal tail of eIF3c (Fig. 4C). The interaction between eIF3c and eIF3-binding domain of eIF4G was previously predicted by sitespecific cross-linking3. The structure shows that this interaction occurs through the same eIF3-binding domain of eIF4G that is involved in the interaction with eIF3l (Fig. 4C and G).

Previously, eIF4F was observed to interact with the 43S complex through the eIF3 subunits -e, -k and -l.^4^ However, these subunits are not present in S. cerevisiae eIF3, and eIF3k and -l are dispensable in *Neurospora crassa* and *Caenorhabditis elegans*.36,37 Thus, these previously identified interactions could not occur in all eukaryotes, raising the question of how eIF4F makes an interaction with the 43S in those species. In this higher-resolution structure, we see several additional interactions between eIF4F and the 43S complex that are likely to be universally conserved. One of these could be the eIF4G-eIF3c interaction, given that the eIF3c C-terminal tail includes residues highly conserved among eukaryotes, including in S. cerevisiae (Fig. S9), which also explain why a knockdown of eIF3c reduces the recruitment of the 43S.38

The eIF4A component of eIF4F binds to a pocket formed by eIF3c, -e, -h, and -l (Fig. 4). We previously described interactions between eIF4A and eIF3e, and eIF3k/l.^4^ The improved local resolution of this post-scanning complex allowed us to further characterize the interaction between eIF4A and the octameric structural core of eIF3. Of the two RecA domains of eIF4A, the C-terminal domain (CTD) is in close proximity to eIF3l, while the NTD makes contact with eIF3c, -e, and -h.

The interactions between eIF4A and eIF3h raises many questions. eIF3h is known to interact with METTL3 and promote mRNA circularization during m6A dependent translation.^39^The two helices from eIF3h extend toward the solvent site of the 43S, but the role of eIF3h in initiation was not clear. The core of the mammalian eIF3 octameric structural core is formed by a 7-helix bundle consisting of two helices from eIF3h and one helix from -c, -e, -f, -l and -k.4,40–43 Here, we see a direct interaction between eIF3h and eIF4A-NTD (Fig. 4I and Fig. S10) through which eIF3h may play a regulatory role in initiation (Fig. 4). Consistent with this idea, phosphorylation at residue Ser183 of eIF3h, located just above the domain that interacts with eIF4A (Fig. S10), has been implicated in the process of reinitiation in plants.^44^. In humans, the same phosphorylation has been associated with a possible oncogenic role of eIF3h.^45^ It is possible that this post-translational modification of eIF3h affects its interaction with eIF4F. Furthermore, this domain of eIF3h also interacts with mRNA (Fig. S10).

Unexpectedly, the structure reveals a direct interaction between eIF4A-NTD and ribosomal protein eS7 (Fig. 4J). The structure rationalizes previous studies on translational control whereby eS7 is monoubiquitinated by the human E3 ubiquitin ligase CNOT4 or its yeast ortholog Not4.46–50 Deubiquitination is required to allow the cap-binding complex eIF4F to bind to the 43S.49,51 Here, we see a direct interaction between eIF4A-NTD and the domain of eS7 ubiquitinated by E3 ubiquitin ligase (Fig. 4J-K). Superimposing our structure with a structure of ubiquitin (Ub)^34^ at the predicted binding site in eS7^51^ places the Ub in a position where it could clash with eIF4A (Fig. 4K), thereby preventing the binding of the cap-binding complex eIF4F, showing why deubiquitination of eS7 is required for translation initiation.49,51

### Upon start-codon recognition, human eIF5-NTD occupies the site vacated by eIF1

Around the site normally occupied by eIF1, there is additional density that cannot be accounted for by the factor (Figs. 1, Fig. S11). Both previous biochemical data and a structure of a yeast initiation complex showed that the yeast N-terminal domain of eIF5 (eIF5-NTD) replaces eIF1 upon start codon recognition.52,53 Rigidbody fitting of the known structure of human eIF5-NTD (PDB: 2E9H) accounted for the entire density present at the platform of the 40S (Fig. S11). Moreover, we could fit the zinc-binding domain (ZBD) of eIF5-NTD into the density (Fig. S11). This domain is not present in eIF1, confirming that this density arises from bound eIF5-NTD that has displaced eIF1 upon start-codon recognition just as was previously proposed for the yeast system.52,53

The structure therefore reveals an evolutionarily conserved set of events occur on startcodon recognition, whose fidelity is enhanced by the factors eIF1, eIF1A and eIF5.54 First, eIF1A and eIF1 bind to the 40S platform near the A and P sites, respectively, and promote the open conformation of the 40S that facilitates the binding to the tRNAiMet. During scanning, the tRNAiMet is not fully inserted into the P site of the ribosome^4^4, which allows codon-anticodon sampling until recognition of the AUG start codon.4,55 After start-codon recognition (Fig. 5A), the tRNAiMet is inserted fully into the P site of the ribosome,40,41,55–59 resulting in a clash with eIF1. Thus, upon start-codon recognition, eIF1 dissociates from the complex and is replaced by eIF5-NTD (Fig. 5). Like eIF1, eIF5-NTD monitors the codon-anticodon interaction (Fig. 5B-C). However, while eIF1 residue Asn39 clashes with the anticodon stem-loop, the eIF5-NTD residue Asn30 adopts a different conformation (Fig. 5D-E), which allows it to monitor codon-anticodon interaction without clashing with the tRNAiMet (Fig. 5C). During scanning, eIF1 Loop 2 interacts with the tRNAiMet D loop, which prevents the accommodation of the tRNAiMet into the PIN state. Upon start codon selection, Loop 2 undergoes conformational changes to allow the accommodation of the tRNAiMet in the PIN state.55,60 The destabilization of eIF1 and its replacement with eIF5-NTD avoids a clash with the tRNAiMet and stabilizes it in the PIN state because eIF5-NTD Loop 2 is both shorter and adopts a different conformation than that of eIF1 (Fig. 5D).

**Figure 5:**
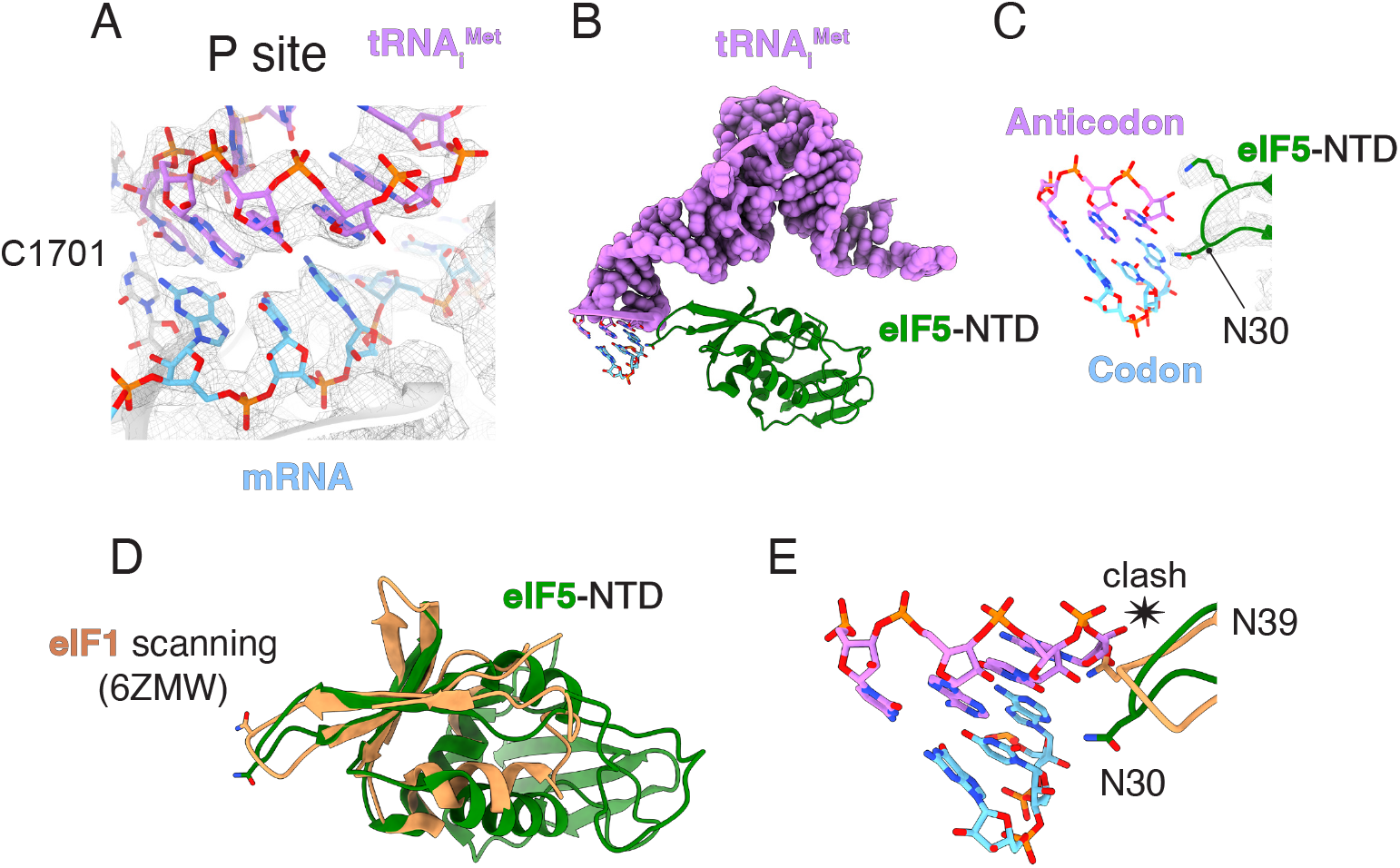
Codon-anticodon base pairing in the P site. (A) Close-up of the P site to highlight the codonanticodon base pairing. (A to C) eIF5-NTD replaces eIF1 after AUG start codon recognition. (D to E) Superposition of eIF5-NTD with the structure of human eIF1 during scanning (PDB:6ZMW).^4^ The conformation of eIF1 during scanning would clash with the anticodon stem loop of the tRNA_i_^Met^ after start codon recognition.

### Molecular mechanism for translational regulation by eIF4A and eIF4F

In our structure, the cap-binding complex eIF4F binds upstream of the 43S complex, while a second molecule of eIF4A binds downstream, at the entry site of the mRNA binding channel. These locations suggest a model where eIF4A plays eIF4F-dependent and eIF4F-independent roles during mRNA recruitment15 and scanning (Fig. 6A-C).

**Figure 6:**
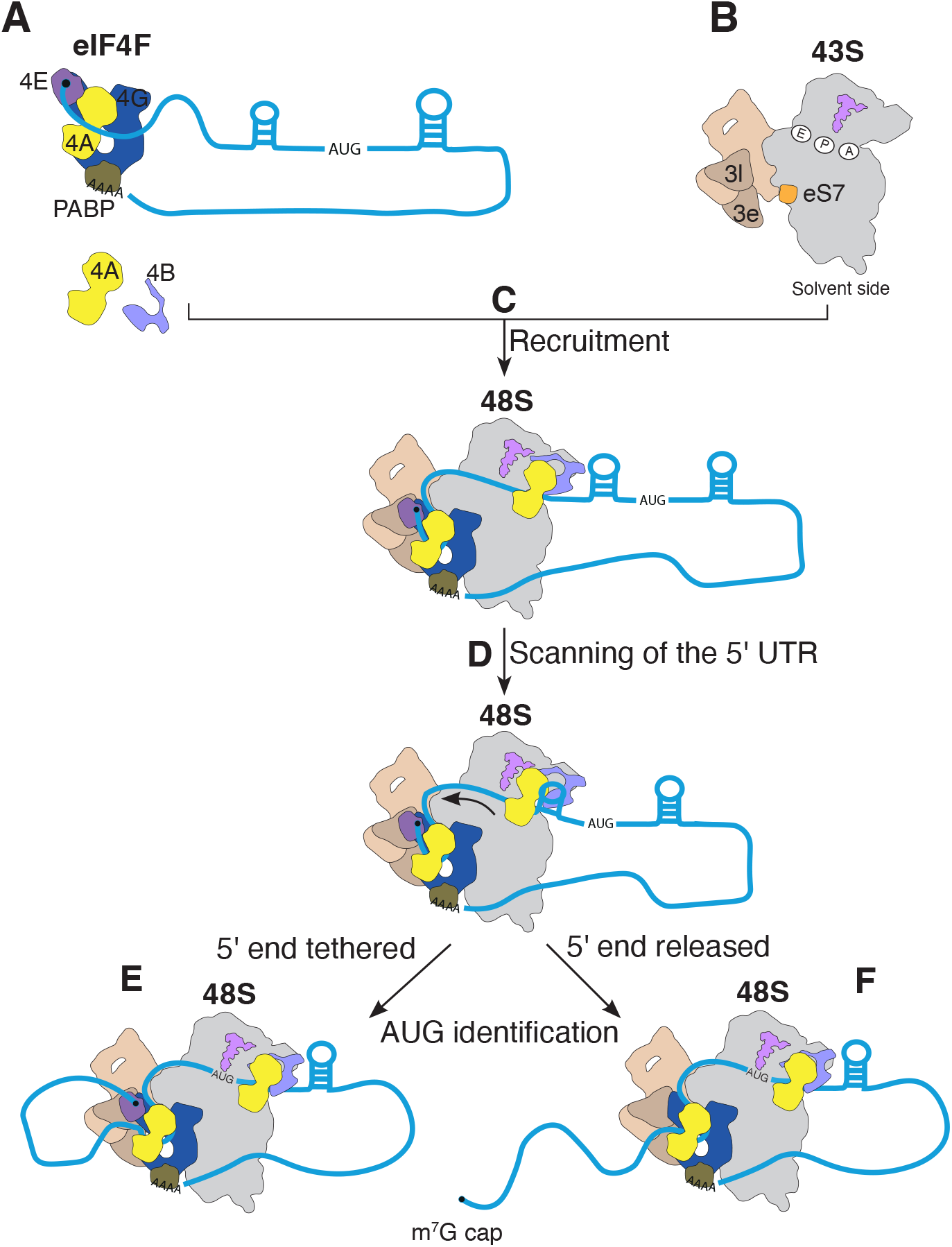
Model for translational regulation by eIF4F and eIF4A. (A to C) The cap-binding complex eIF4F (A) binds at the 5^′^end of mRNA and recruits the 43S (B) to form the 48S complex (C). eIF4F binds to eIF3 and ribosomal protein eS7 located upstream of the 43S. The position of eIF4F is compatible with a model whereby the mRNA is slotted into the mRNA channel in the 40S small ribosomal subunit. In addition to eIF4F, a second molecule of eIF4A at the mRNA entry site is likely to play a role in this process by facilitating the accommodation of the mRNA into the channel. The ATPase activity of eIF4A is stimulated by eIF4B and eIF3bgi module^15^ located at the entry site. Thus, eIF4A at the entry site is likely to use the energy from ATP hydrolysis to unwind the RNA secondary structure downstream of the 43S (D). It remains unclear whether eIF4E remains attached to the rest of eIF4F (E) or is released (F) during the scanning process, but the structure suggeststhelatter

The cap-binding complex eIF4F at the 5′ end of mRNA plays a critical role in recruiting the 43S downstream of it, in a process that involves the mRNA being slotted into its channel in the 40S subunit (Fig. 6C). The higher resolution of the current 48S complex reveals how eIF4F makes universally conserved interactions with the 48S complex, including eIF3c and ribosomal protein eS7.

Canonical eIF4F-dependent initiation is enhanced by a topologically closed loop of the mRNA, often mediated by an interaction between eIF4G at the 5′ end and PABP at the 3′ end.61,62 Such a topology would require a slotting mechanism for recruitment. Similarly, during m6A-dependent translation, it has been shown that the mRNA is circularized through an interaction between METTL3 located at the 3′ of mRNA and eIF3h.39 Other modes of initiation also require the slotting mechanism, including that via internal ribosome entry sites (IRES), eIF3d-dependent or circular mRNA. However, some mRNAs with unusually short 5′ UTR (less than 40 nucleotides in mammals) may use alternative recruitment pathways. A recent study proposed that these mRNAs may use the slotting mechanism as well.63 However, this would require a backward movement of the 43S (3′ to 5′) rather than the small oscillations that that are known to exist.22 Thus, it is more likely that those transcripts use alternative recruitment pathway. In addition to eIF4F, a second eIF4A helicase is coordinated by the 40S subunit and eIF3 so that it binds to mRNA before it enters the mRNA binding channel (Fig. 6C). The function of this entry site bound eIF4A may promote the accommodation of the mRNA into the entry channel, especially when secondary structure is present in the 5′ UTR.15,64 Consistent with this idea, the cryo-EM 3D classification reveals a class of particles with eIF4A bound to the 43S but without the mRNA accommodated into the channel at the entry site (Fig. S12), suggesting that the binding of the second eIF4A to the 43S precedes the binding of the mRNA. Given the possible location of the eIF4B-RRM in our structure, it is possible that eIF4B may also play a role in the mRNA accommodation process.^65^

The position of eIF4A at the entry site of the mRNA binding channel provides strong evidence that this conserved RNA helicase likely functions to unwind the mRNA secondary structure as it enters the mRNA binding channel during scanning. Thus, we propose a model whereby after recruitment, both eIF4A and eIF4F work in concert to ensure the directionality of scanning (Fig. 6D). In this model, eIF4F would prevent the reverse movement of the 43S complex^4^ while the second molecule of eIF4A at the leading edge of the ribosome would use the energy from ATP hydrolysis to unwind downstream RNA secondary structure and translocate along the 5′ UTR of mRNA during scanning (Fig. 6D). It is possible that ATP hydrolysis by eIF4F might further help the movement of the 43S and help overcome energy barriers encountered by the entry channel eIF4A.

Compared to many other helicases, eIF4A has low helicase activity. Even though its ATPase activity is stimulated by the 43S and its co-factor eIF4B, scanning of 5′ UTR of mRNA with highly stable secondary structure may require additional helicases, such as mammalian DHX29, which has been seen to bind at the mRNA channel entry site.42,66 In our structure, eIF4A-eIF4B interacts with the 43S at the exact location that DHX29 does (Fig. S13). Thus, it is possible that for initiation on specific mRNAs with high secondary structure, other helicases such as DHX29 are recruited to the 43S complex instead of a second eIF4A molecule. Such a method would allow additional control of translation initiation of such mRNAs.

eIF4E stabilizes the binding of mRNA to eIF4G-eIF4A^33^ and stimulates the helicase activity of the eIF4F bound eIF4A.67 Selective 40S ribosome profiling has indicated that scanning is cap-tethered in most human cells^68^ (Fig. 6E). In the prior structure of 48S there is a low-resolution density that we tentatively assigned to eIF4E.^4^ That structure used a very short mRNA, so no significant scanning had occurred. In the current structure, scanning has spanned 105 nucleotides before the start codon was reached, thus representing a later stage of initiation. Although local resolution was significantly improved, we did not see such additional density that we could assign to eIF4E, suggesting that it, along with the 5′ cap, is released from eIF4F and the rest of the 48S during scanning (Fig. 6F).

In conclusion, our structural data reveal how two eIF4A helicase proteins coordinate the binding of mRNA to the entry and exit sites of 40S subunit mRNA binding channel. Importantly, the entry channel bound eIF4A helicase is precisely positioned to enable its ATPase activity to directly unwind secondary structure located downstream of the scanning 48S complex. Our suggested location of eIF4B adjacent to this second eIF4A molecule would explain its role in the helicase activity of eIF4A. The eIF4A helicase has become an important therapeutic target, with eIF4A-binding natural products showing promising anti-tumor activity in pre-clinical studies. The discovery that two copies of eIF4A are bound to the 48S complex may therefore help guide strategies for cancer therapy.

## Methods

### Purification of human eIFs

Ribosome, tRNAiMet, and human eIFs, including eIF4G557-1105, were purified as described previously.59,64,67,69 PABP was purified as described.^67^ Additionally, we used a hydroxyapatite column to remove any remaining nucleic acids bound to it, and the protein was purified further by gel filtration chromatography on a Superdex 200 Increase column in a buffer containing 20 mM HEPES, pH7.5, 200 mM KCl, 10% glycerol, 1 mM TCEP.

### *In vitro* transcription and purification of mRNA

We designed a synthetic DNA containing a T7 promoter to generate an mRNA with a 5′ UTR containing three polypurine motifs followed by an AUG codon (underlined):

GGACAAGAGAGAGAGAGACUCCAACUCCAAGAGAGAGAGA-GACAACUCCAAGAGAGAGAGAGACAAACCCTCGCUGAGCC-GCAGUCAGAUCCUAGCGUCGAGUUGAUGCUGUCCGAU.

The construct was purchased as a gene block from IDT. After linearization, the DNA plasmid was used as a template for in vitro transcription. The mRNA was purified using a denaturant acrylamide gel followed by electroelution. The purified mRNA was capped using Vaccinia Capping System (New England Biolabs) and then polyadenylated (∼90 nucleotides) (Fig. S1) using E. coli Poly(A) polymerase (New England Biolabs). After free nucleotide, enzyme, and buffer removal, the pure mRNA was stored in water. We used the same protocol of transcription and purification of DLP mRNA used for cryo-EM:

GACACUUGCUUUUGACACAACUGUGUUUACUUGCAAUCCC-CCAAAACAGACAGAAUGAACAACGAGCCACCGCAAACCAG-UGACGGCCGCCGGAGGCGCCCGCGCCCGGCGGCCGAGAGA-GCAGAACGAGACCACACGGAUCCGAGAAGAUUCAUCCUCC-UUCAAUGCCUGGAGGAUA

### *In vitro* reconstitution of human 48S complexes for cryo-EM

We reconstituted the 48S complex by mixing the 43S with eIF4F, eIF4A, eIF4B, PABP, and mRNA in a 25 μl reaction. To reconstitute the 43S, we mixed 0.5 μM 40S with 0.9 μM eIFs and 1.8 μM TC to a final volume of 18 μl in a buffer (97 mM KAc, 2.5 mM MgAc, 3% glycerol, 0.1 mM spermidine, 1mM DTT and 0.5 mM GMP-PNP). We incubated the 43S reaction mix at 30 °C for 10 min. In parallel, we assembled the cap-binding complex eIF4F by mixing a copurified eIF4G (residues 165-1599)-eIF4A complex with eIF4E, mRNA, and PABP, to a final concentration of 3.3µM in 12µl reaction in a buffer (97 mM KAc, 2.5 mM MgAc, 3% glycerol, 0.1 mM spermidine, 1mM DTT and 0.5 mM ATP-g-S).Finally, we mixed 1 μM eIF4F, 1 μM eIF4B, 1 μM eIF4A, 43S (0.3 μM 40S, 0.5 μM eIFs), and 0.5 mM Rocaglamide A in a 25 μl reaction. The reaction mix was incubated at 30 °C for 10 min. The same protocol was used to assemble the 48S complexes with eIF4G_557-1105_, without eIF4B, and with DLP mRNA. The DLP 48S was assembled without Rocaglamide A.

### Cryo-EM grid preparation

The assembled 48S complexes were crosslinked using 1.5 mM BS3 (final concentration) on ice for 45 min to prevent dissociation of eIFs during the grid preparation. As described before,^4^ the crosslink reaction in the presence of spermidine (a polyamine with quenching properties) and on ice has a mild effect — comparable with the complex without BS3. 3 μl of 140 nM 48S complex (based on the concentration of the 40S) was applied onto UltrAuFoil R1.2/1.3 300 mesh gold grids pre-covered with graphene oxide (Sigma) suspension made in-house. Briefly, the UltrAuFoil gold grids were washed using deionized water and then dried overnight at room temperature. The grids were then glow-discharged for 5 min at 30 mA, followed by incubation with 3 μl of graphene oxide (0.2 mg/ml) for 1 min. After the incubation and blotting, the grids were washed three times using 20 μl deionized water. The grids were dried for at least 1h before at room temperature and then used. We prepared the grids using an FEI Vitrobot Mark IV at 4 °C and 100% humidity. The grids were blotted for 7 or 8 sec at blotting force -15 and then plunged into liquid ethane at 93 K in a precision cryostat system produced at the MRC LMB.70

### Cryo-EM data collection

Data were collected on Titan Krios microscopes (ThermoFisher) equipped with a K3 direct electron detector camera (Gatan) at a magnification of 105,000x and at pixel sizes of 0.826 Å pix-1 or 0.829 Å pix-1 (48S data set from eBIC/Diamond). The data were collected using EPU software, super-resolution counting mode using a Bioquantum energy filter (Gatan) (binning 2), faster acquisition mode, and with defocus ranging from -1.2 μm to -3.0 μm. Doses/frames: 1.197 e/Å^2^ (48S data set from eBIC/Diamond); 1.2391 e/Å^2^ (48S data set from MRC LMB); 0.6889 e/Å^2^ (48S without eIF4B and 48S complexes with eIF4G557-1105). DLP 48S was collected using Falcon 4 electron detector, counting mode and 1 e/Å^2^/frame.

### Image processing

Motion correction was performed using the implementation in RELION 4.71 Movies were aligned using 5 □ 5 patches with dose-weighting. CTF was estimated using CTFFIND4.1.^72^ After 2D classification, we used the cryo-EM map of a human 48S^4^ after lowpass filtering to 60 Å as a reference for 3D classification. After 3D classification and refinement, we performed mask classification to select only particles containing eIF3 octameric structural core. We used Bayesian polishing in RELION to correct beam-induced motion.71 After polishing, we performed mask classification at the entry site, followed by the eIF4F binding region and TC. Finally, we used multi-body refinement and flexibility analysis in RELION^73^ to account for the predominant molecular motions.

### Model building, fitting, and refinement

We used our previous structure of human 48S,^4^ previous cryo-EM and crystal structures of eIFs^17,23,25,26,40,42,52,58,74–76^, as well as AlphaFold,^35^ to build and refine the atomic model. Fitting and model building and local refinement were performed in Coot.77 We used Phenix for real space refinement.78

## Figures

All figures were made using ChimeraX.79

## Data availability

The atomic model and cryo-EM maps have been uploaded to the Protein Data Bank (PDB:XXXX) and to the Electron Microscopy Data Bank (EMD:YYYY).

**Figure S1:**
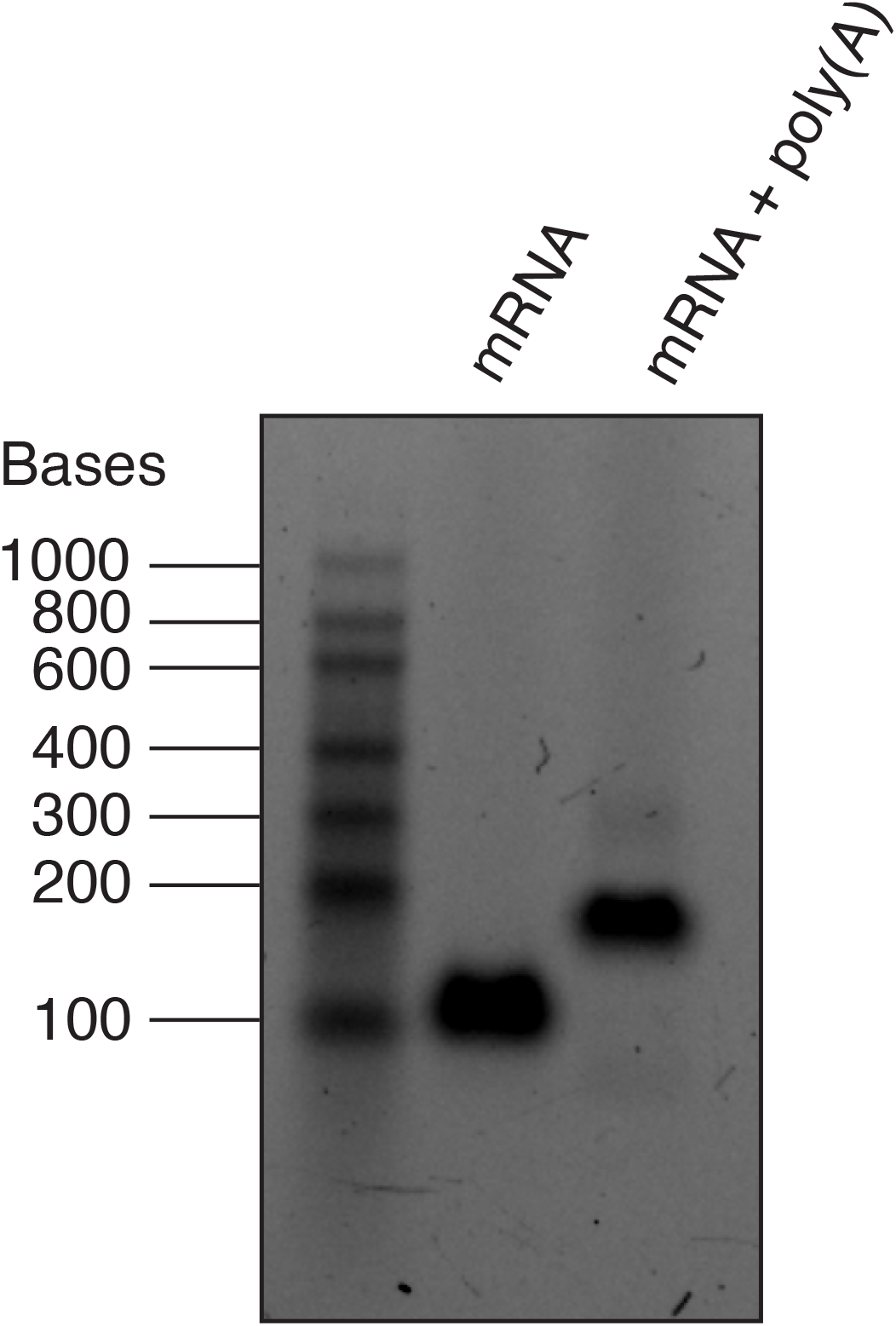
Analysis of capped mRNA after Poly(A) tailing. The polyadenylated mRNA has additional 90

**Figure S2:**
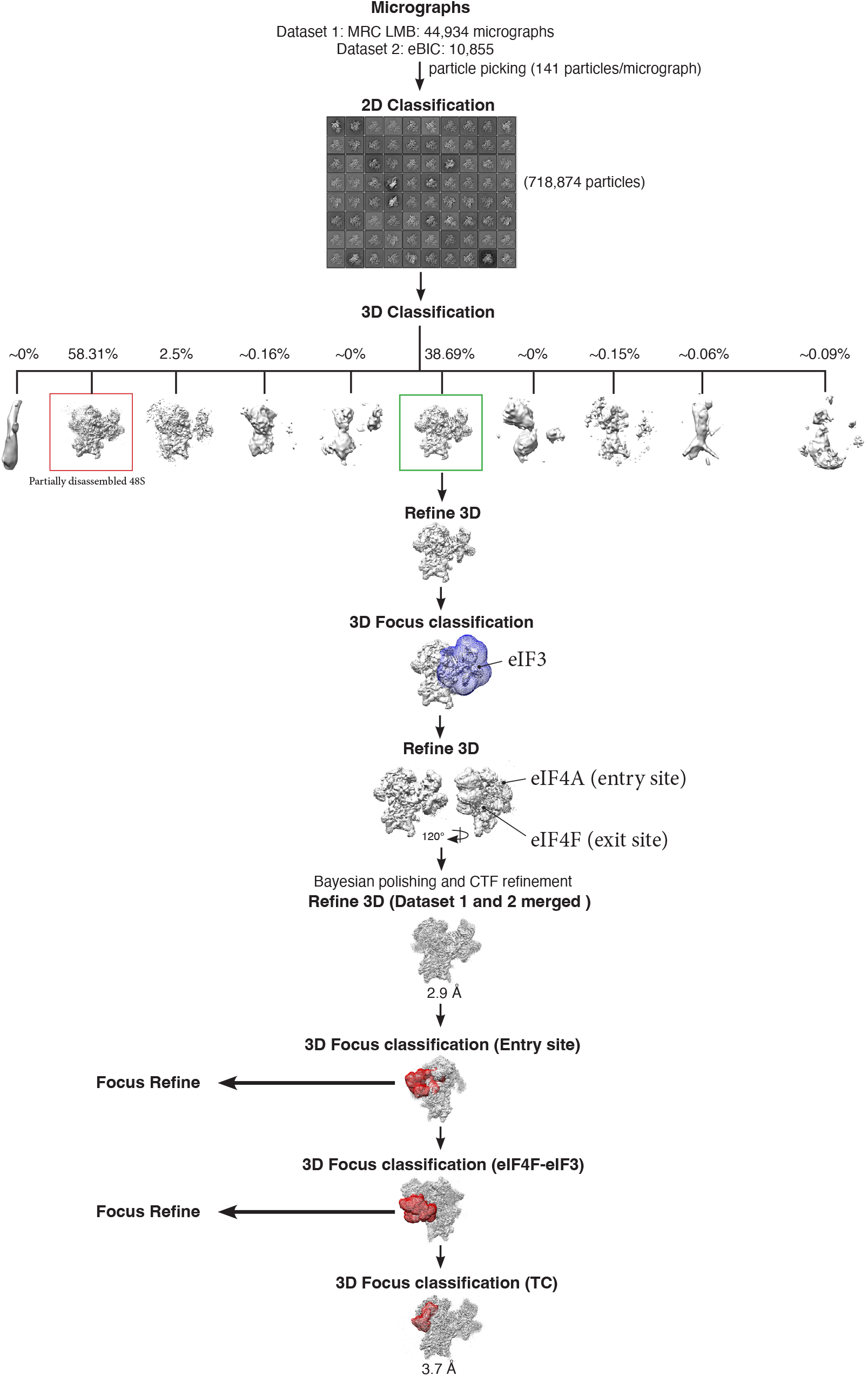
Cryo-EM data analysis of human 48S. Datasets 1 and 2 were analyzed independently. After polishing and CTF refinement, the two datasets were merged, which yielded the 2.9 Å 3D reconstruction of a human 48S.

**Figure S3:**
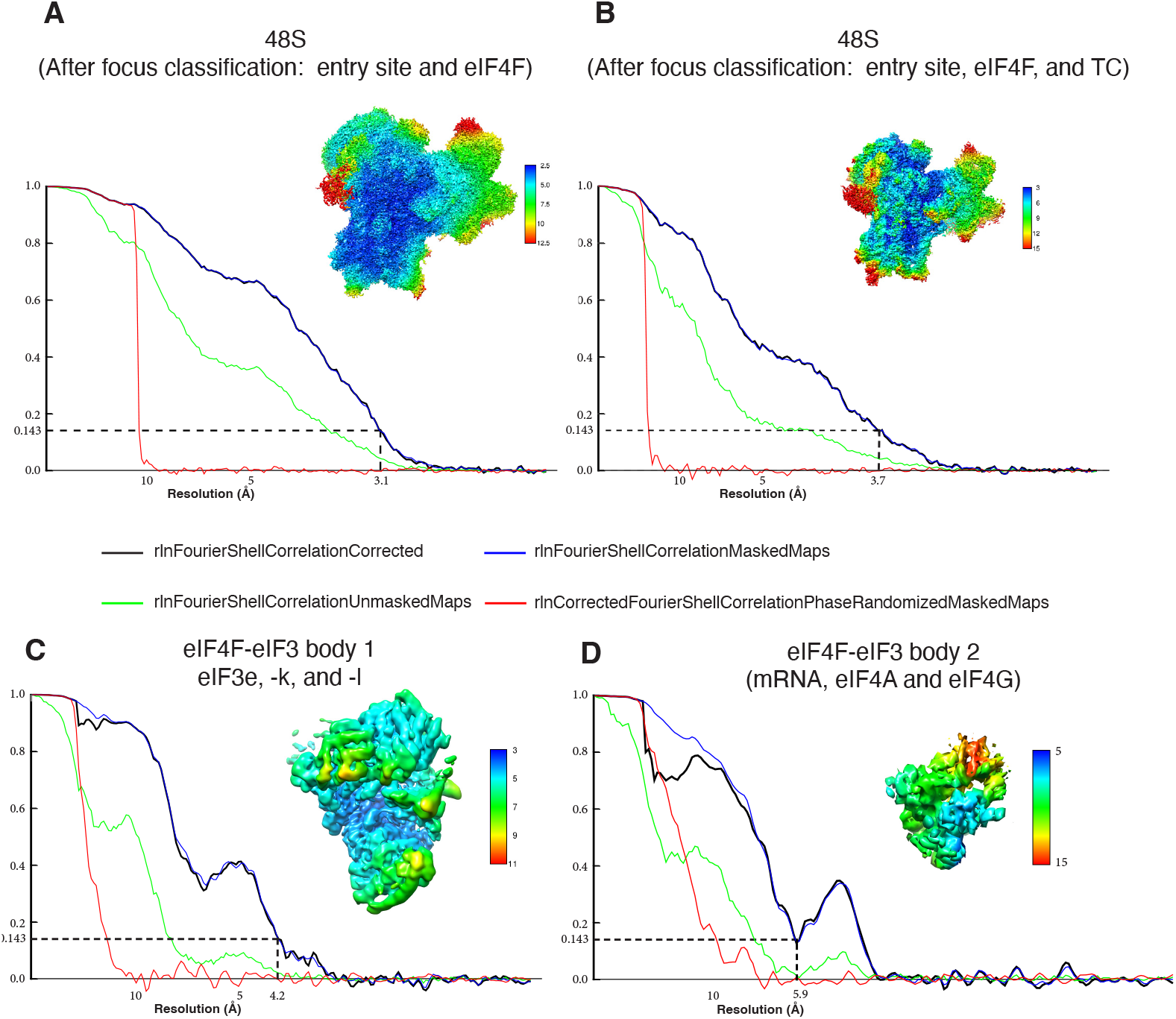
Overall and local resolution. (A and B) Fourier shell correlation (FSC) curve and local resolution of the 48S complex after focus 3D focus classification at the entry site, eIF4F, and ternary complex. (C and D) Overall and local resolution of eIF4F-eIF3 after multi-body refinement.

**Figure S4:**
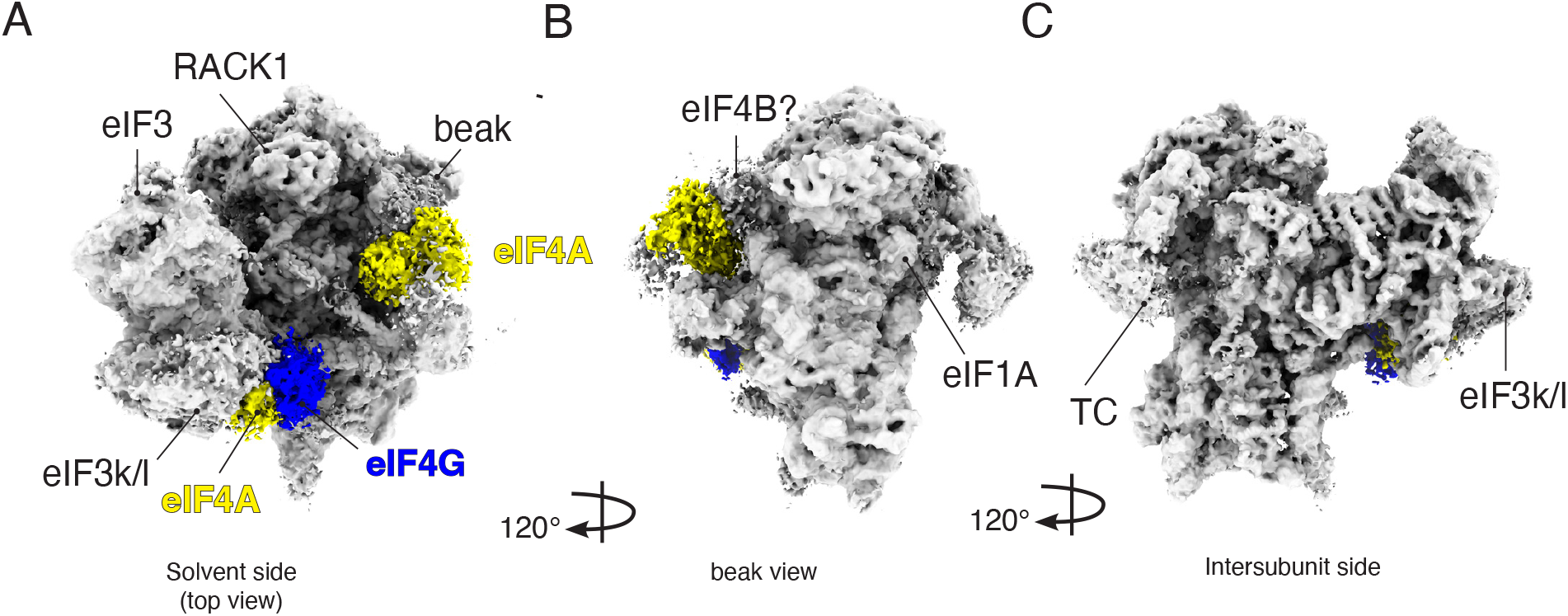
Cryo-EM 3D reconstruction of a human 48S complex assembled without Rocaglamide (RocA). (A to C) Overview of the cryo-EM density map. The complex contains both eIF4F and the second molecule of eIF4A bound at the mRNA channel entry site. To assemble this complex, we used an mRNA with a highly stable RNA stem-loop structure (DLP) located 83 nucleotides downstream of the cap. This mRNA also has an AUG start codon.

**Figure S5:**
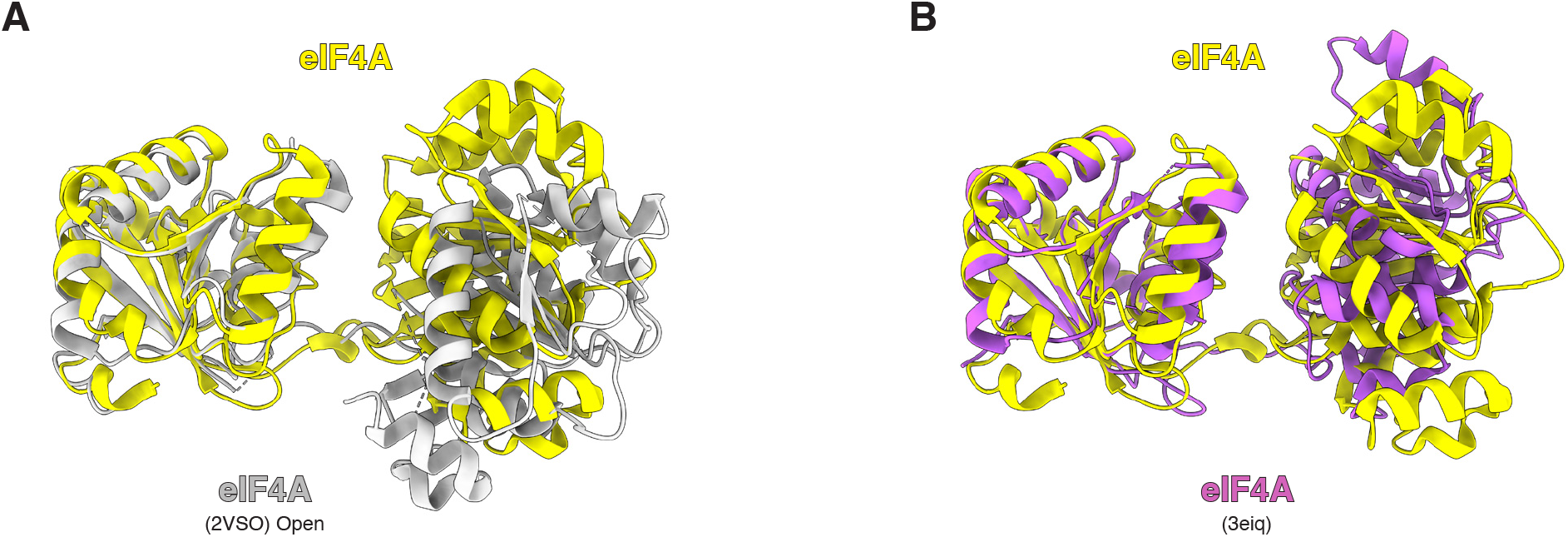
Conformation of molecule of eIF4A bound at the mRNA entry site. (A) Superposition of eIF4A with the structure of a yeast eIF4A in complex with eIF4G (PDB:2VSO)^23^. (B) Superposition of eIF4A with the structures of eIF4A in complex with Pdcd4 (PDB:3eiq)^24^.

**Figure S6:**
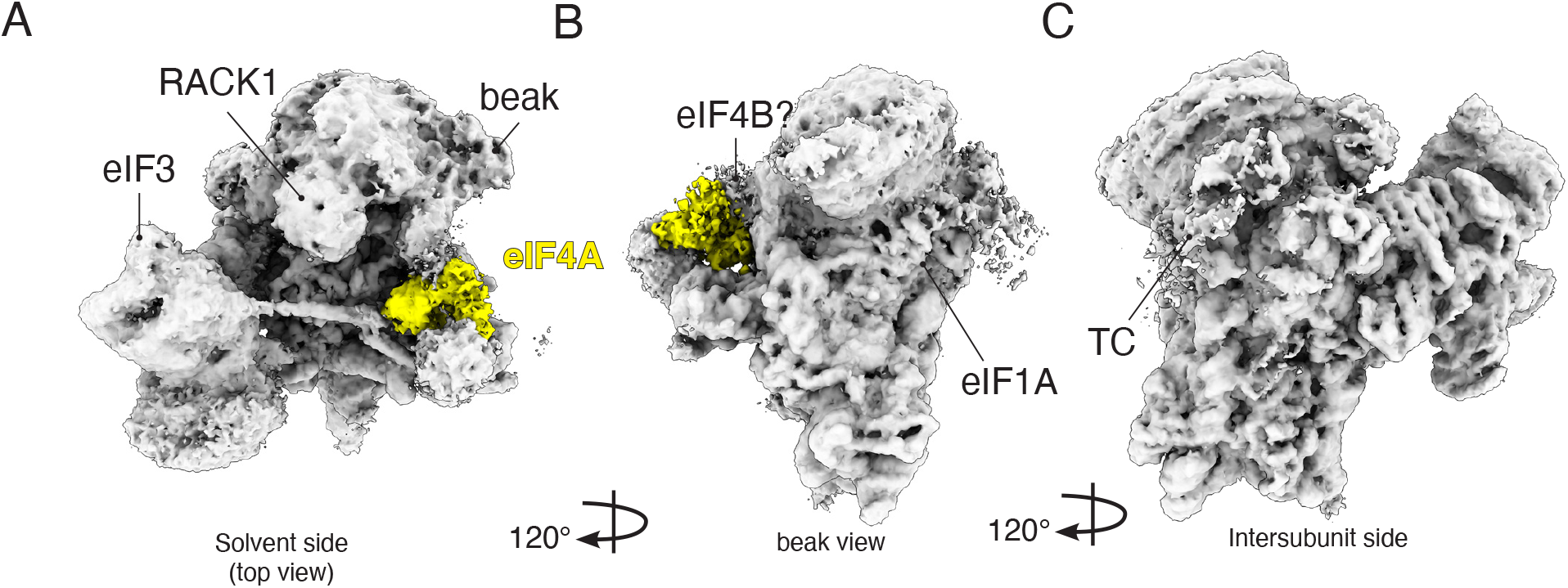
Cryo-EM 3D reconstruction of a human 48S assembled with a truncated eIF4G (residues 557-1136). (A to C) Overview of the cryo-EM density map. The eIF4A density is visible even without 3D focus classification.

**Figure S7:**
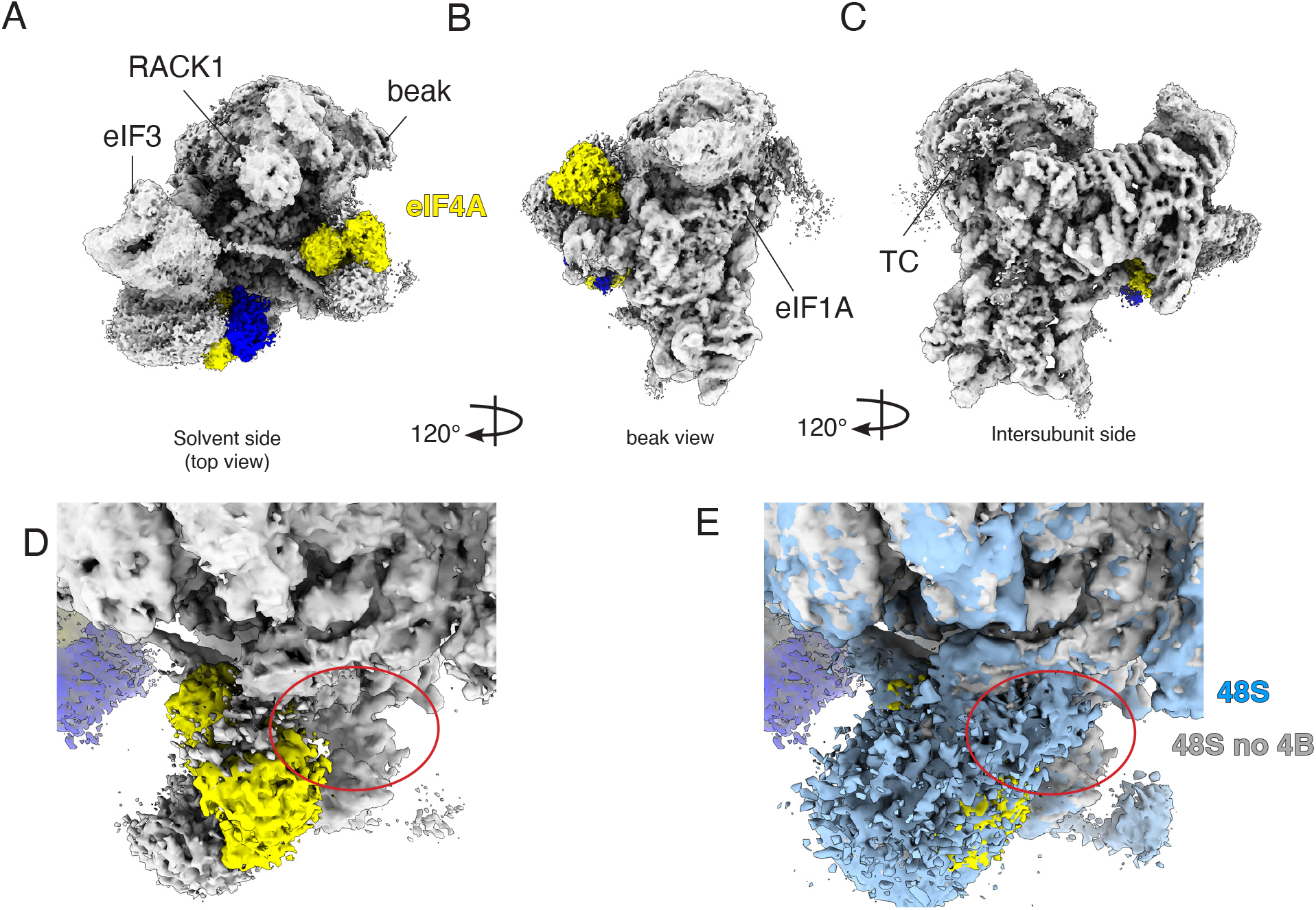
Cryo-EM 3D reconstruction of a human 48S assembled without eIF4B. (A to C) Overview of the cryo-EM density map. The red oval highlights the absence of the density in the complex without eIF4B (D) that can be seen in the structure of 48S assembled eIF4B (E).

**Figure S8:**
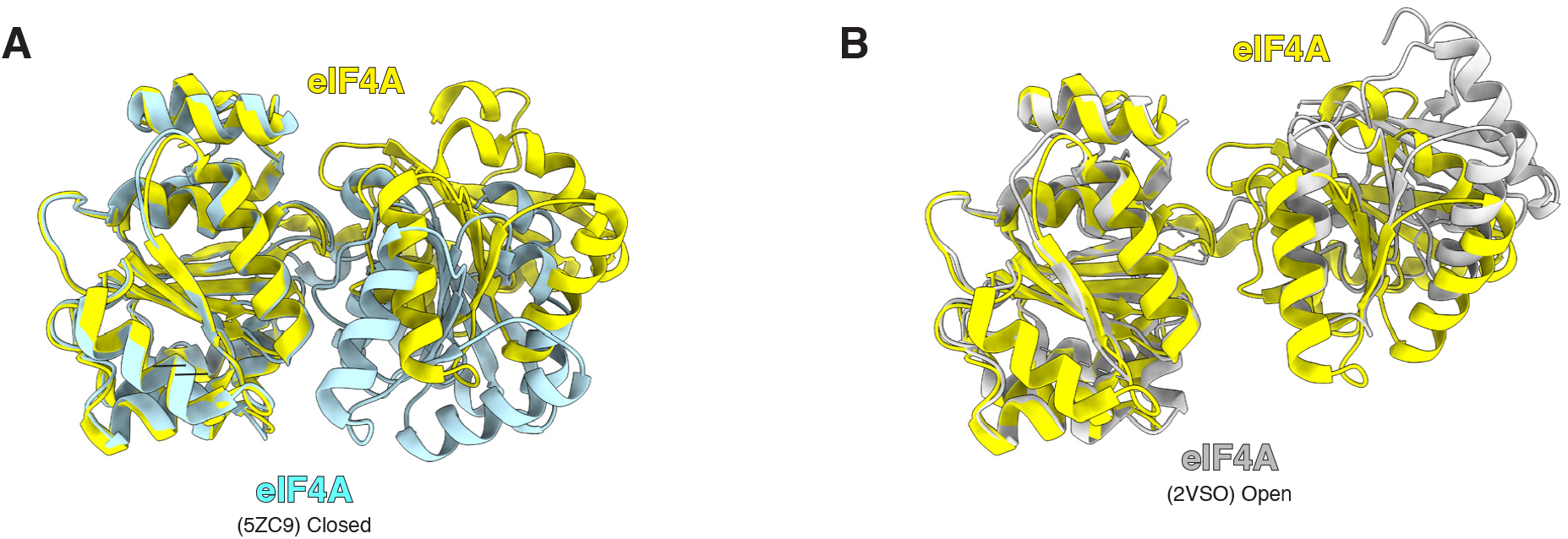
Conformations of the eIF4A in the cap-binding eIF4F complex. (A) Superposition of eIF4A that is part of the cap-binding eIF4F complex in the 48S with the structures of human free eIF4A in complex with mRNA and RocA (PDB:5ZC9)^17^ (B) Superposition of this eIF4A with the structures of a yeast eIF4A in complex with eIF4G (PDB:2VSO)^23^ *Homo sapiens Oryctolagus cuniculus Saccharomyces cerevisiaeDrosophila melanogaster*

**Figure S9:**
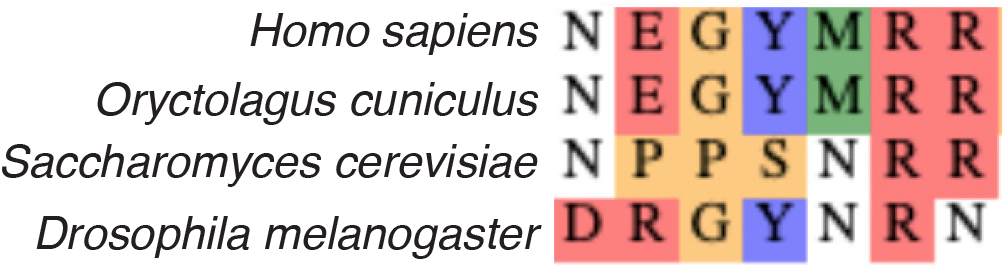
Multiple sequence alignment of the eIF3C C-terminal tail. The site of interaction with eIF4G is conserved among eukaryotes.

**Figure S10:**
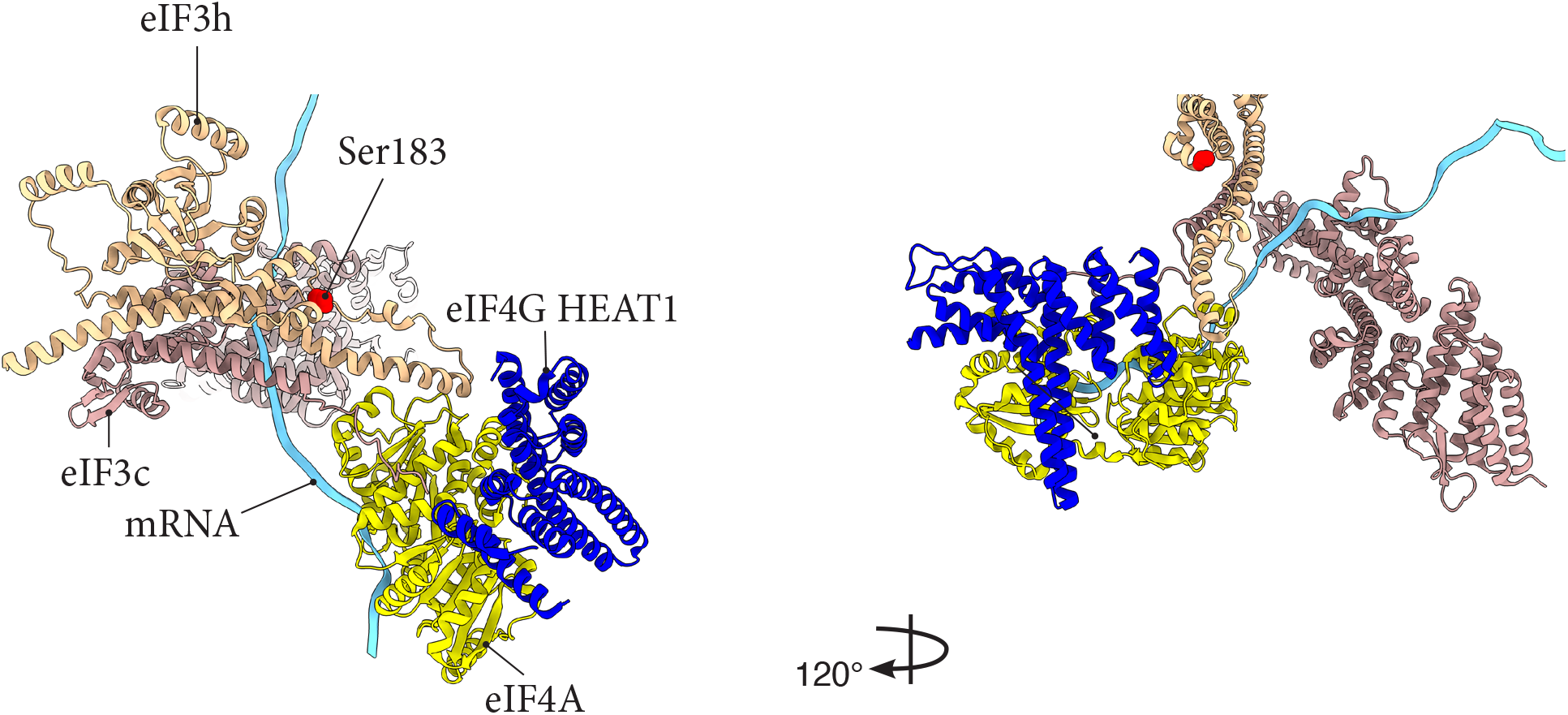
Interaction of eIF3h and eIF3c with eIF4F. eIF3h interacts directly with eIF4A-NTD. eIF3h and eIF3c provide clamping and guide the mRNA from eIF4F towards the mRNA channel in the 40S.

**Figure S11:**
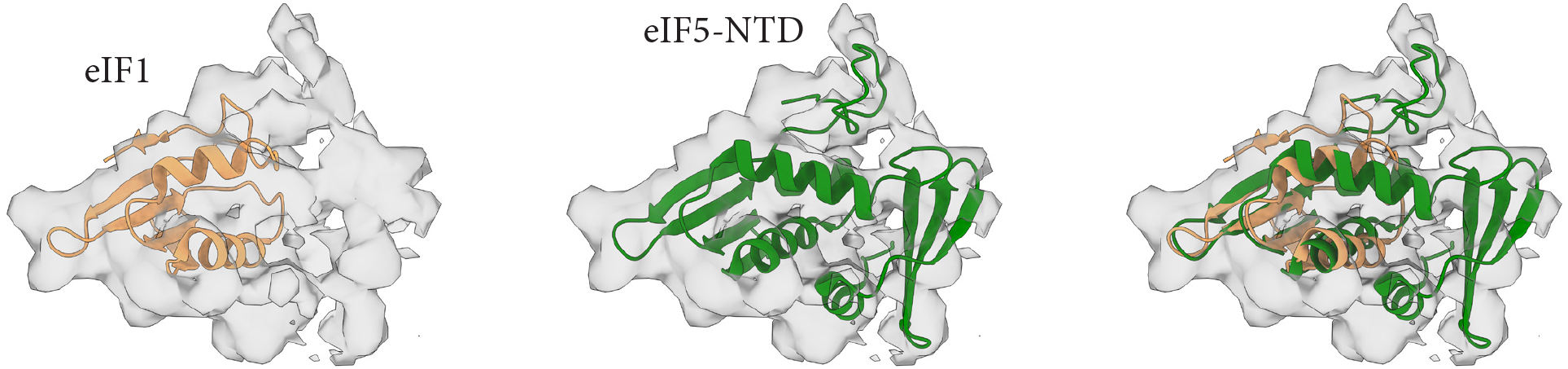
Map-to-model fits of human eIF1 and eIF5. Rigid-body fitting of human eIF1 (PDB:6ZMW)^4^ and eIF5-NTD (PDB: 2E9H) into the cryo-EM map.

**Figure S12:**
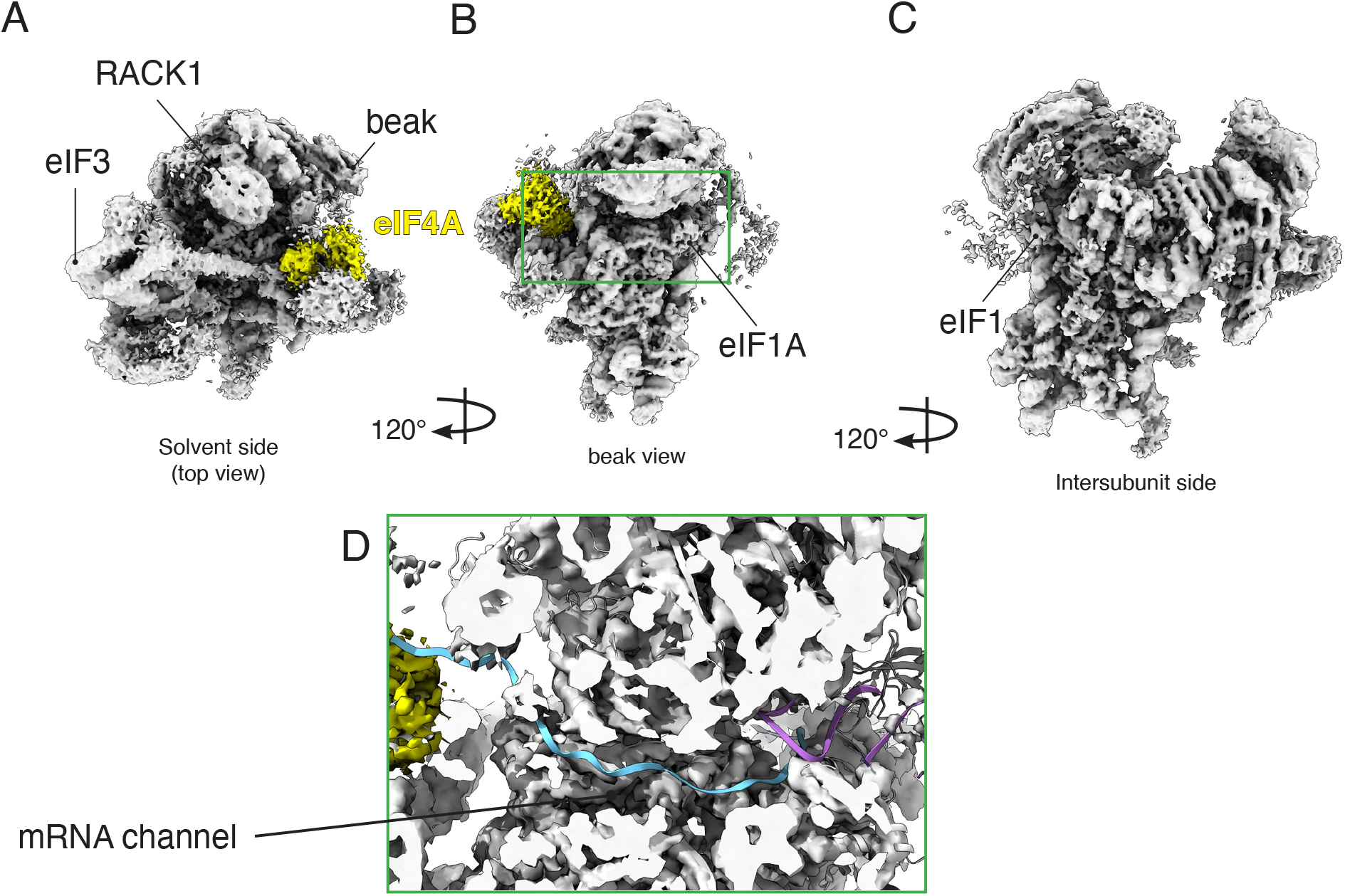
Cryo-EM 3D reconstruction of a class of particles without mRNA into the channel. (A to C) Overview of the cryo-EM density map. (D) Inset showing lack of density for mRNA using mRNA and tRNA superimposed from the 48S structure.

**Figure S13:**
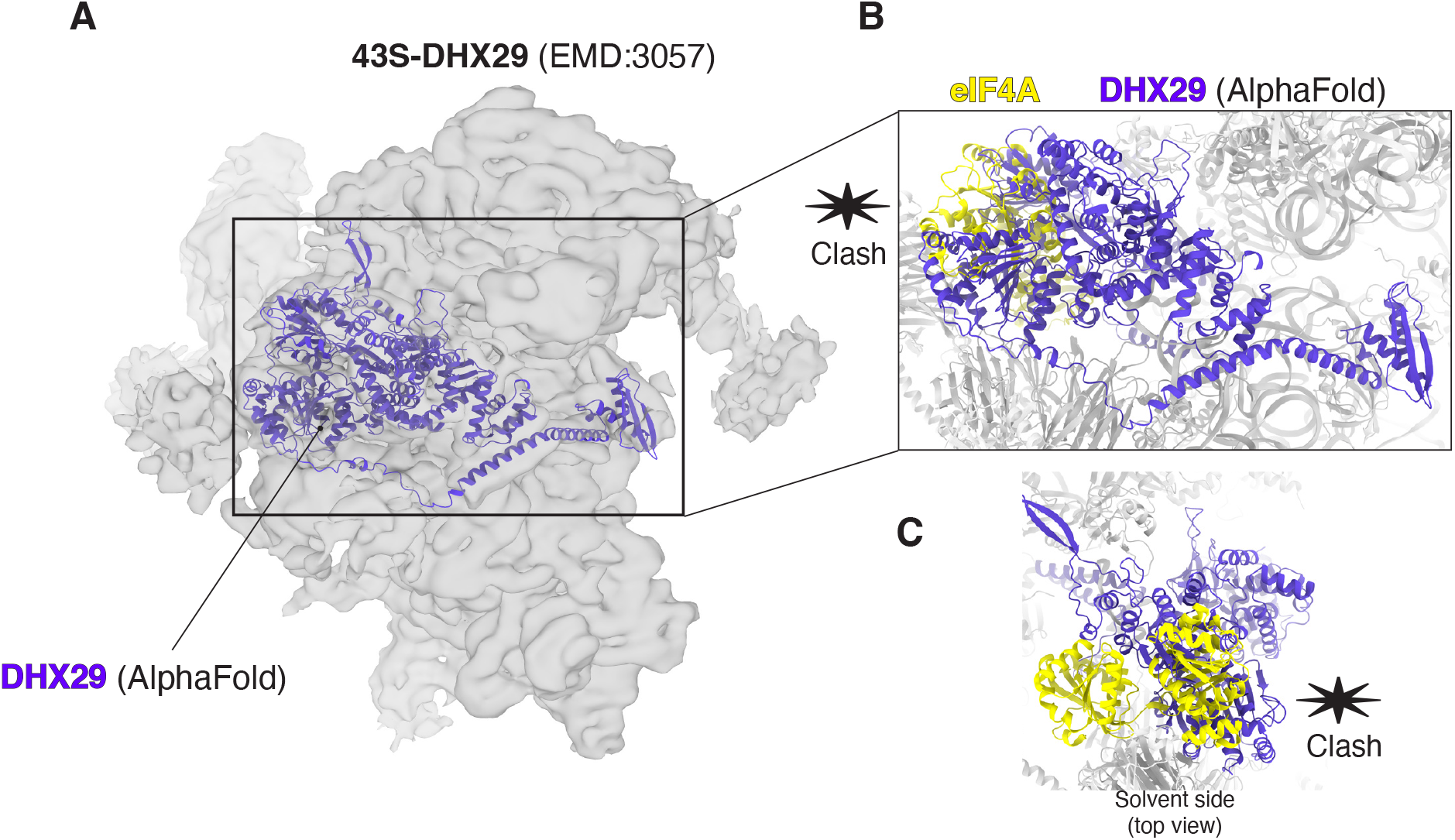
eIF4A interacts with the 43S in the same location as DHX29. (A) Rigid-body fitting of human DHX29 (AlphaFold prediction) into a prior cryo-EM map of a rabbit 43S (EMD:3057)^42^. (B and C) The inset shows that DHX29 and the entry-site eIF4A would overlap and could not coexist.

**Table S1.**
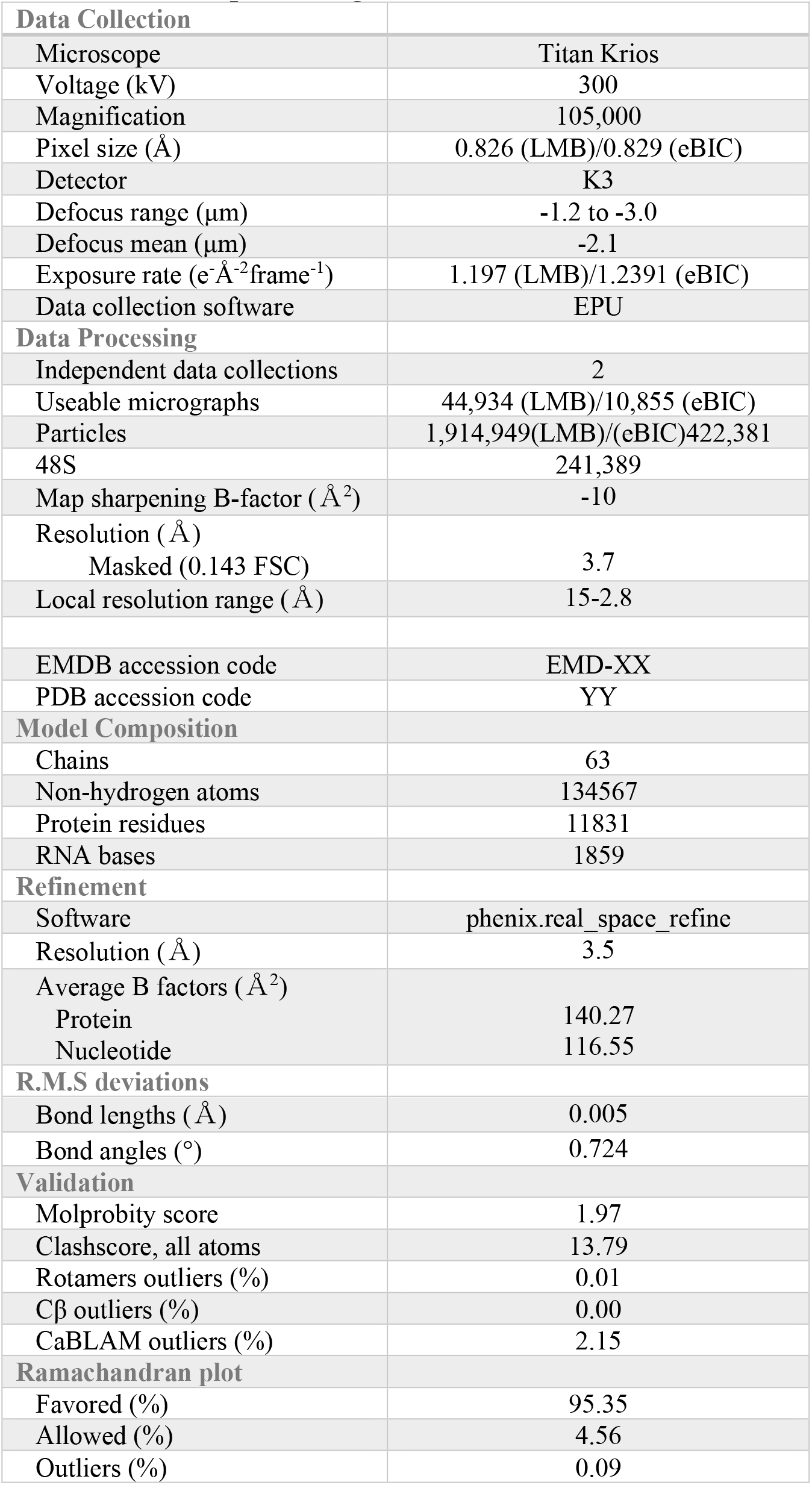
Data collection, processing, refinement and model statistics.

